# Poly(A) tail length is a major regulator of maternal gene expression during the mammalian oocyte-to-embryo transition

**DOI:** 10.1101/2021.08.29.458052

**Authors:** Yusheng Liu, Hu Nie, Chuanxin Zhang, Zhenzhen Hou, Jiaqiang Wang, Falong Lu

## Abstract

Transcription is silent during the mammalian oocyte-to-embryo transition (OET) until zygotic genome activation (ZGA). Therefore, the OET relies on post-transcriptional regulation of maternal mRNA, among which poly(A) tail lengths have been found to regulate translation for a small number of genes^1–3^. However, transcriptome-wide poly(A) tail length dynamics and their role in gene expression during the mammalian OET remain unknown. Here, we quantified transcriptome-wide mRNA poly(A) tail length dynamics during the mammalian OET using PAIso-seq1 and PAIso-seq2^4,5^, two methods with different underlying principles that preserve the poly(A) tail information. We revealed that poly(A) tail length was highly dynamic during the mouse OET, and Btg4 is responsible for global maternal mRNA deadenylation. We found that the poly(A) tail length positively associated with translational efficiency transcriptome-wide in mouse oocytes. In addition, genes with different alternative polyadenylation isoforms show longer poly(A) tails for isoforms with distal polyadenylation sites compared to those with proximal polyadenylation sites in mouse, rat, pig and human oocytes after meiotic resumption, which is not seen in cultured cell lines. Surprisingly, mammalian embryos, namely mouse, rat, pig, and human embryos, all experience highly conserved global mRNA re-polyadenylation after fertilization, providing molecular evidence that the early embryo development before ZGA is driven by re-polyadenylated maternal mRNAs rather than newly transcribed mRNAs. Together, our study reveals the conserved mRNA poly(A) tail length landscape. This resource can be used for exploring spatiotemporal post-transcriptional regulation throughout the mammalian OET.

## Introduction

The non-templated poly(A) tail is an essential part of most mature mRNA transcripts, and regulates their translation efficiency (TE) and stability^6–13^. Poly(A) tail length is controlled by two opposite processes: polyadenylation and deadenylation. Post-transcriptional regulation through modulating the length of poly(A) tails is known to functionally impact biological processes including gamete development, tumor metastasis, innate immunity, inflammation, obesity, synaptic plasticity, higher cognitive function, and long-term memory^13–24^.

The oocyte-to-embryo transition (OET) in mammals refers to the developmental processes occurring between meiotic resumption of oocytes arrested at prophase of meiosis I and zygotic genome activation (ZGA) after fertilization, and it is essential for successful reproduction of the next generation^1–3^. One of the distinctive features of the OET is the silencing of transcription prior to ZGA^13,25^. Given the lack of active transcription, the diverse biological events occurring during the OET, which in addition to ZGA include meiotic progression, fertilization, zygote activation, pronuclei formation, epigenome reprogramming, and totipotency establishment^26–34^, are largely controlled by the population of maternal mRNAs that is stored in oocytes. These mRNAs are tightly regulated at the post-transcriptional level to ensure reproduction success^1,3^.

Although poly(A) tails are thought to be a major determinant of the fate of maternal mRNAs, there is to date no record of the global dynamics of the mRNA poly(A) tails during the mammalian OET. TAIL-seq (or the modified version, mTAIL-seq) and PAL-seq are two technologies developed based on the Illumina platform that have helped to greatly deepen our understanding of the regulation of poly(A) tail length in various organisms, but these technologies have not been used to study the mammalian OET because of the requirement for a large amount of input materials and the limited availability of mammalian oocytes and embryos^18,35,36^.

To explore the transcriptome-wide regulation of mRNA poly(A) tails, we previously developed two leading accurate methods in sequencing mRNA poly(A) tails named PAIso-seq1 (previously named PAIso-seq) and PAIso-seq2^4,5^. Here, using these two methods, we were able to reveal the global mRNA poly(A) tail length dynamics at different stages during the mouse OET, namely the GV (germinal vesicle), GVBD (GV breakdown), MI (metaphase I), and MII (metaphase II) stages for oocytes and the 1C (1-cell), 2C (2-cell), and 4C (4-cell) stages for embryos. We revealed the transcriptome-wide association of poly(A) tail length and translational efficiency in maturing mouse oocytes. We provide transcriptome-wide molecular evidence for deadenylation and degradation of maternal mRNA through Btg4. Interestingly, alternative polyadenylation (APA) isoforms with a distal polyadenylation site (dPAS) have longer poly(A) tail length than those with a proximal polyadenylation site (pPAS) in maturing oocytes, which is conserved in mice, rats, pigs, and humans. Surprisingly, transcriptome-wide global re-polyadenylation is seen in all four of these species after fertilization. Thus, our study reveals the conserved spatiotemporal post-transcriptional regulation of mRNA poly(A) tail length during mammalian OET.

## Results

### Transcriptome-wide poly(A) tail length dynamics during the mouse OET

To globally measure poly(A) tail length dynamics during the OET, we applied the PAIso-seq2 and PAIso-seq1 methods to analyze mouse GV, GVBD, MI, and MII stage oocytes and 1C, 2C, and 4C stage embryos. Very interestingly, we observed a highly dynamic pattern of global poly(A) tail length (Fig. 1a). The PAIso-seq2 data revealed that the poly(A) tail length distributions of the GV and GVBD stages are similar to that seen in somatic cells^35,36^, whereas the poly(A) tail length distribution of the MII stage is highly biased toward very short lengths (Fig. 1a). The poly(A) tails in the stages before (MI) and after (1C) the MII stage are longer than those in the MII stage (Fig. 1a). The highly dynamic pattern of poly(A) tail length distribution was also seen in the PAIso-seq1 dataset (Extended Data Fig. 1a), confirming the highly dynamic nature of global poly(A) tail length during the mouse OET. Next, we looked into the change in poly(A) tail length of individual genes between consecutive stages (Fig. 1b). There was minimal global change in poly(A) tail length between the GVBD and GV stages, whereas drastic changes were observed between the MI vs GVBD, MII vs MI, 1C vs MII, 2C vs 1C, and 4C vs 2C stages (Fig. 1b). As transcription is silent before ZGA during the OET (GV, GVBD, MI, MII, and 1C stages), the poly(A) tail length changes we detected represent post-transcriptional regulation of the poly(A) tails. In particular, the shift of global poly(A) tail length toward longer tails from the MII to 1C stage caught our interest, and is described in more detail in a later section.

**Fig. 1.**
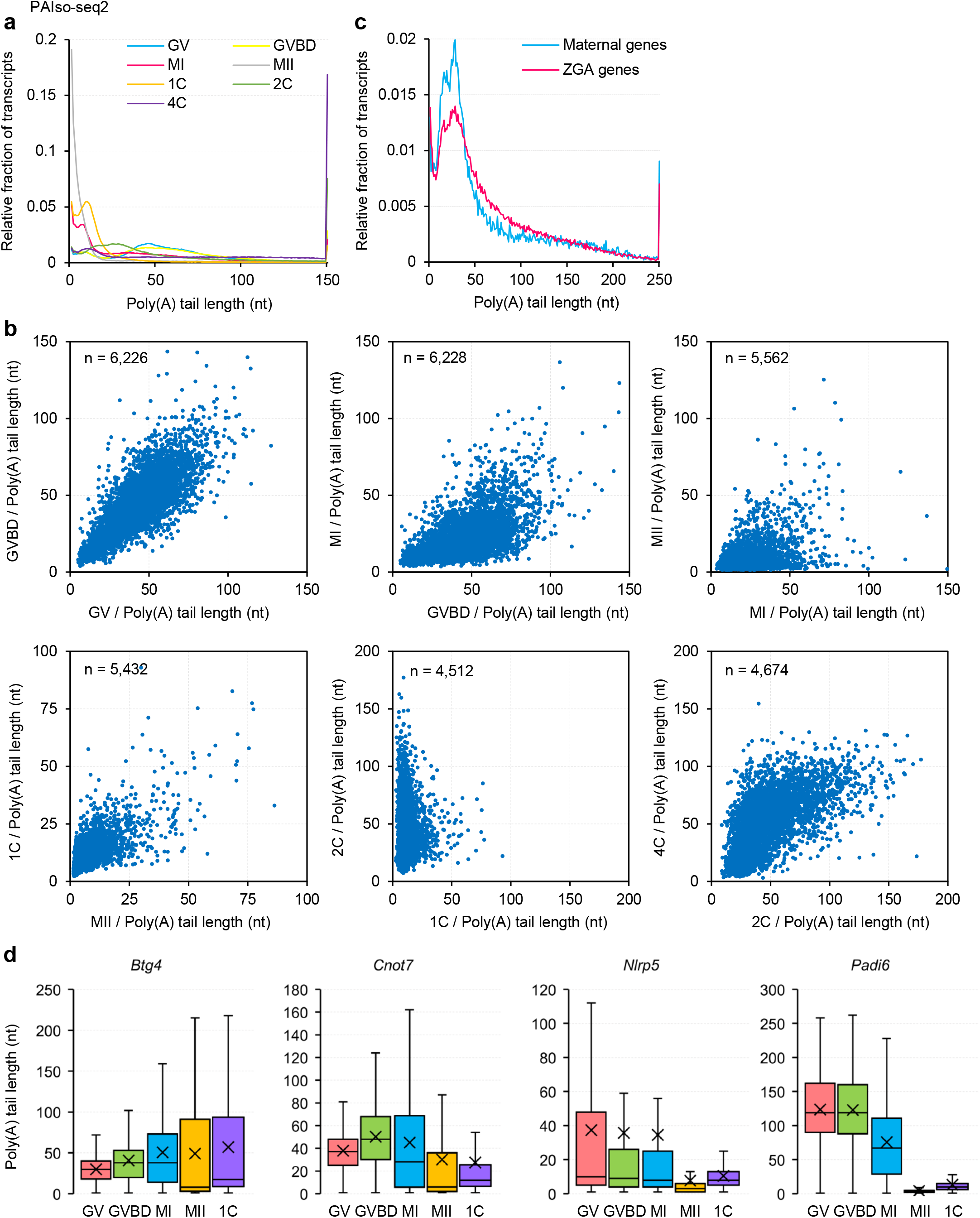
Dynamics of mRNA poly(A) tail length during the mouse OET. **a,** Histogram of transcriptome-wide poly(A) tail lengths in mouse oocytes and early embryos measured by PAIso-seq2. Histograms (bin size = 1 nt) are normalized to cover the same area. Transcripts with a poly(A) tail of at least 1 nt are included in the analysis. Transcripts with a poly(A) tail length greater than 150 nt are included in the 150 nt bin. **b,** Scatter plots of poly(A) tail length of mouse samples at neighboring developmental stages. Each dot represents one gene. The poly(A) tail length for each gene is the geometric mean length of all the transcripts with a poly(A) tail of at least 1 nt for the given gene. For each plot, genes with at least 10 reads in both of the samples are included in the analysis. The number of genes included in the analysis is shown on the top left of each plot. **c,** Histogram of poly(A) tail lengths of combined transcripts from maternal genes and zygotic genes in 2-cell stage mouse embryos. Histograms (bin size = 1 nt) are normalized to cover the same area. Transcripts with a poly(A) tail of at least 1 nt are included in the analysis. Transcripts with a poly(A) tail length greater than 250 nt are included in the 250 nt bin. **d,** Box plot of the poly(A) tail lengths of *Btg4*, *Cnot7*, *Nlrp5*, and *Padi6* in mouse samples at different stages. The “×” indicates mean value. The black horizontal bars show the median value. The top and bottom of the box represent the 25^th^ and 75^th^ percentiles, respectively. Transcripts with a poly(A) tail of at least 1 nt are included in the analysis.

The burst of ZGA gene transcription is known to occur at the 2C stage in mouse^37^. Therefore, we looked into the 2C stage data, which consist of reads from a mixture of maternal and zygotic transcripts. Indeed, the maternal transcripts in 2C embryos, which are already much shorter than those in GV oocytes, showed obviously shorter poly(A) tails than those of zygotic transcripts produced during ZGA (Fig. 1c and Extended Data Fig. 1b), suggesting that the maternal mRNAs are undergoing deadenylation-mediated decay at the 2-cell stage, while the newly synthesized ZGA transcripts have relatively long poly(A) tails. It is well known that mRNA transcripts of the *Nlrp5* (*Mater*) and *Padi6* maternal genes are robustly translated in GV oocytes; their gene products are essential for the mouse OET^38–40^. We found that, consistent with the observed global trend, the poly(A) tail lengths of these maternal genes gradually decreased as oocytes developed (Fig. 1d). Although the global trend was decreasing poly(A) tail length during oocyte development, there were individual genes with different patterns of change in poly(A) tail length during the OET. For example, the *Btg4* and *Cnot7* mRNAs are known to be expressed, but they stay dormant in GV oocytes before they start to be translated in the MI stage^28–30,41^. We observed that the poly(A) tail lengths of *Btg4* and *Cnot7* transcripts increased between the GV and MI stages, coinciding with the previously reported translational activation of these transcripts (Fig. 1d).

Taken together, our oocyte and embryo data from two different and complementary methods reveal the transcriptome-wide dynamic landscape of mRNA poly(A) tail lengths during the mouse OET, which represents a valuable resource for studying the post-transcriptional regulation of the mouse OET.

### Poly(A) tail length is associated with the translation efficiency in maturing mouse oocytes

Increased mRNA tail length positively regulates the translation of several individual genes in the oocytes of mice^28,30, 41–43^. However, it is not known whether this is a general mechanism in mouse oocytes. Translation within GV, MI, and MII oocytes in mice has been quantified by analyzing the ribosome engagement of mRNAs using RiboTag/RNA-seq^44^. We compared the poly(A) tail lengths from both the PAIso-seq1 and PAIso-seq2 datasets with the TE of individual genes in GV, GVBD, and MI stage oocytes^44^, and observed a clear positive correlation between ribosome engagement and the length of poly(A) tails in all three stages (Fig. 2a, c). Comparing consecutive developmental stages, we found that changes in poly(A) tail length (from both the PAIso-seq1 and PAIso-seq2 datasets) between the different stages were correlated with the changes in ribosome association, although the correlation between the GV and GVBD stages was relatively low, likely due to the relatively small changes in the poly(A) tail length (Fig. 2b, d).

**Fig. 2.**
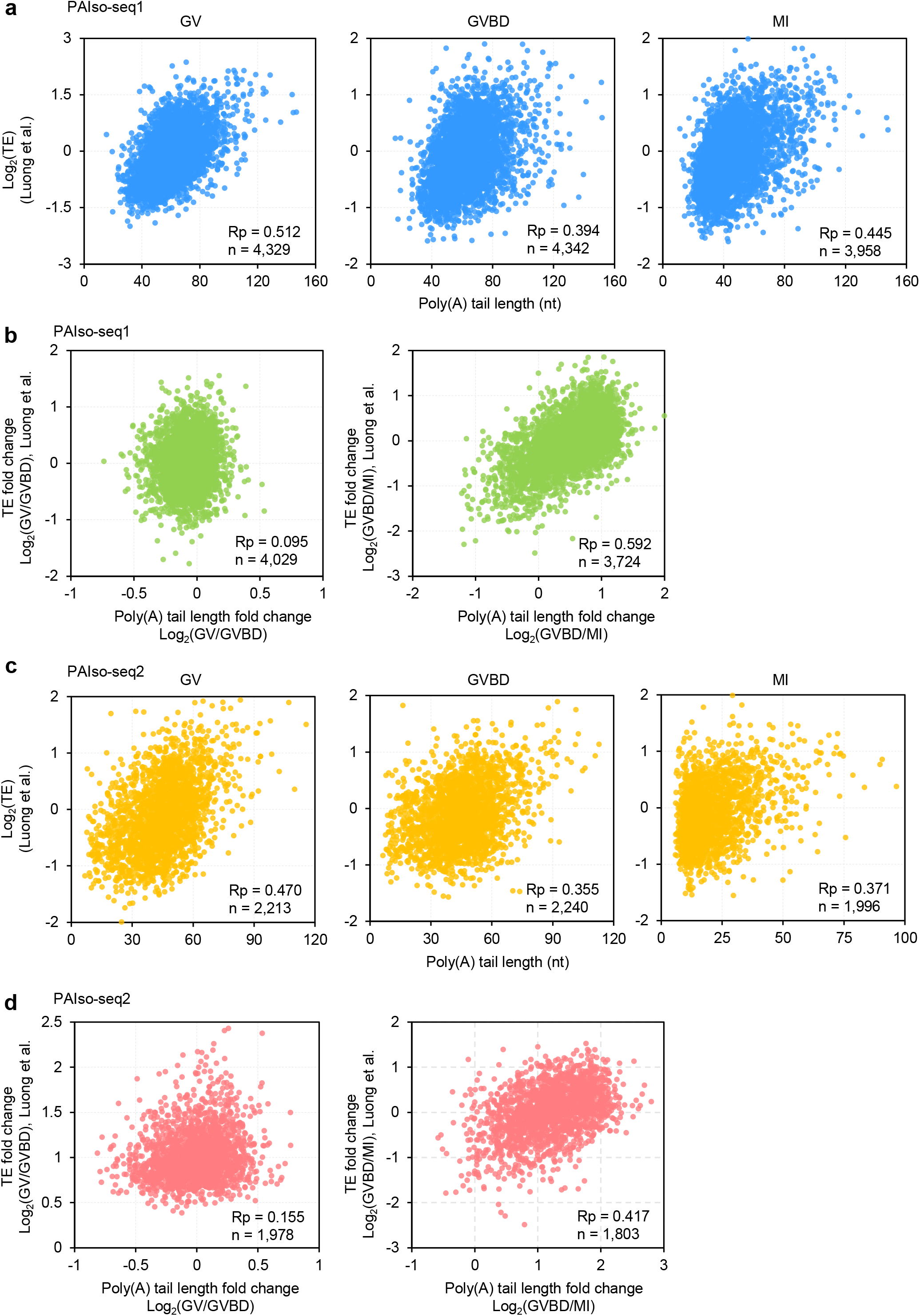
Poly(A) tail length is associated with translational efficiency in maturing mouse oocytes. **a, c,** Scatter plots of poly(A) tail length measured by PAIso-seq1 (**a**) or PAIso-seq2 (**c**) and translational efficiency (TE) of mouse GV, GVBD, and MI oocytes^44^. Each dot represents one gene. The poly(A) tail length for each gene is the geometric mean length of all transcripts with a poly(A) tail of at least 1 nt for the given gene. Genes with at least 20 reads in each of the replicates are included in the analysis. Pearson’s correlation coefficient (Rp) and number of genes included in the analysis are shown on the bottom right. **b, d,** Scatter plots of the change in poly(A) tail length measured by PAIso-seq1 (**b**) or PAIso-seq2 (**d**) and change in translational efficiency (TE) of the neighboring mouse oocyte stage^44^. Each dot represents one gene. The poly(A) tail length for each gene is the geometric mean length of all transcripts with a poly(A) tail of at least 1 nt for the given gene. Genes with at least 20 reads in each of the replicates of both analyzed stages are included in the analysis. Rp and number of genes included in the analysis are shown on the bottom right.

Next, we asked whether genes from the same family are regulated similarly. The *Zp* family proteins are known to be highly abundant in GV oocytes. We found that the poly(A) tail lengths of *Zp1*, *Zp2*, and *Zp3* all gradually decreased from the GV to MI stage, which correlated well with the decrease in TE (Fig. 3a). However, this does not represent a general rule. *Oosp1* is known to be translationally activated at the MI stage^44^. Indeed, we saw that the poly(A) tail lengths of *Oosp1* and *Oosp3* increased from the GV to MI stage (Fig. 3b). However, the change in the poly(A) tail length of *Oosp2* followed an opposite trend (Fig. 3b). The *Cpeb* family genes *Cpeb1*, *Cpeb2*, and *Cpeb3* also showed different trends of poly(A) tail changes (Fig. 3c). In addition, the changes in poly(A) tail length of these genes correlated well with their changes in ribosome association. Therefore, poly(A) tail length-mediated translational regulation is highly gene specific in mouse oocytes, and can be differentially regulated among members of the same gene family.

**Fig. 3.**
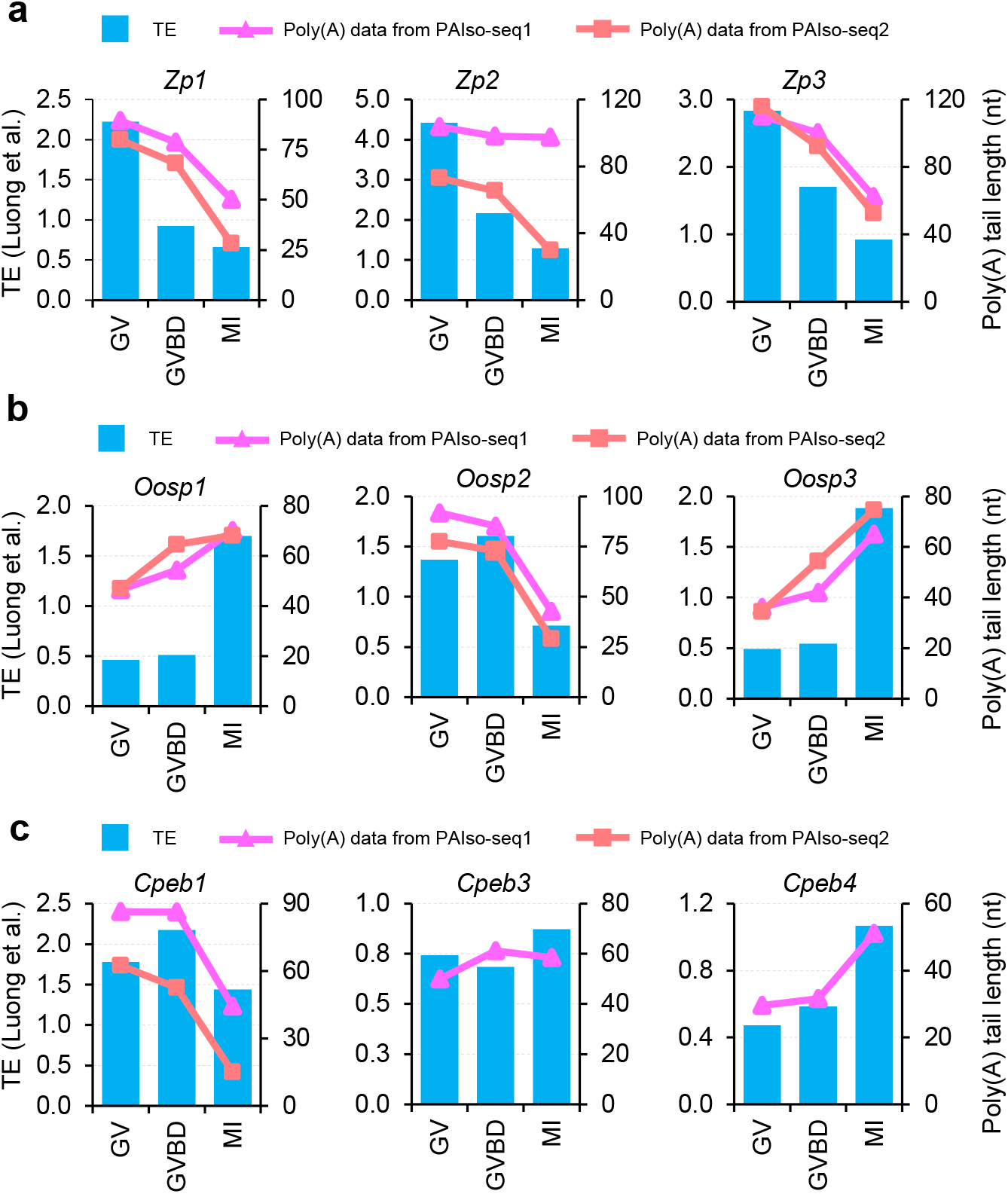
Differential regulation of poly(A) tail length and TE for genes in the same family during mouse oocyte maturation. Plots of poly(A) tail length and translational efficiency^44^ for *Zp* family (**a**), the *Oosp* family (**b**), and the *Cpeb* family (**c**) genes during mouse oocyte maturation. Bar plots show TE, while line plots show the poly(A) tail length measured by PAIso-seq1 or PAIso-seq2. The poly(A) tail length for each gene is the geometric mean length of all transcripts with a poly(A) tail of at least 1 nt for the given gene.

Therefore, our poly(A) tail length data from both PAIso-seq1 and PAIso-seq2 reveal the positive correlation between mRNA poly(A) tail length and TE transcriptome wide during mouse oocyte maturation, suggesting that mRNA poly(A) tail length dynamics shape the highly regulated translational dynamics during mammalian oocyte maturation.

### APA isoforms can be differentially regulated at the level of poly(A) tail length in maturing mouse, rat, pig, and human oocytes

Both the PAIso-seq1 and PAIso-seq2 methods enable full-length mRNA isoform sequence data to be obtained together with poly(A) tail information. Therefore, isoform-specific poly(A) tail length can be analyzed on a transcriptome-wide scale. In general, the usage of a proximal polyadenylation site (pPAS) or distal polyadenylation site (dPAS) is relatively consistent between oocytes at different stages as measured by the ratio of the number of sequenced reads with a dPAS or pPAS for each gene (Fig. 4a, b). Very interestingly, we found that although the poly(A) tail lengths were similar between pPAS and dPAS transcripts in the GV stage, the dPAS transcripts showed relatively longer poly(A) tails compared with pPAS transcripts in GVBD, MI, and MII stage oocytes in mouse (Fig. 4c, d). Furthermore, this finding was conserved in mammalian oocytes. We observed similar differences in poly(A) tail dynamics between the pPAS and dPAS isoforms during oocyte maturation in rats, pigs, and humans (Extended Data Fig. 2). It is likely that the mRNAs undergo global deadenylation in general, but that the cis-elements in the 3’-UTR of the dPAS isoforms protect them from global deadenylation.

**Fig. 4.**
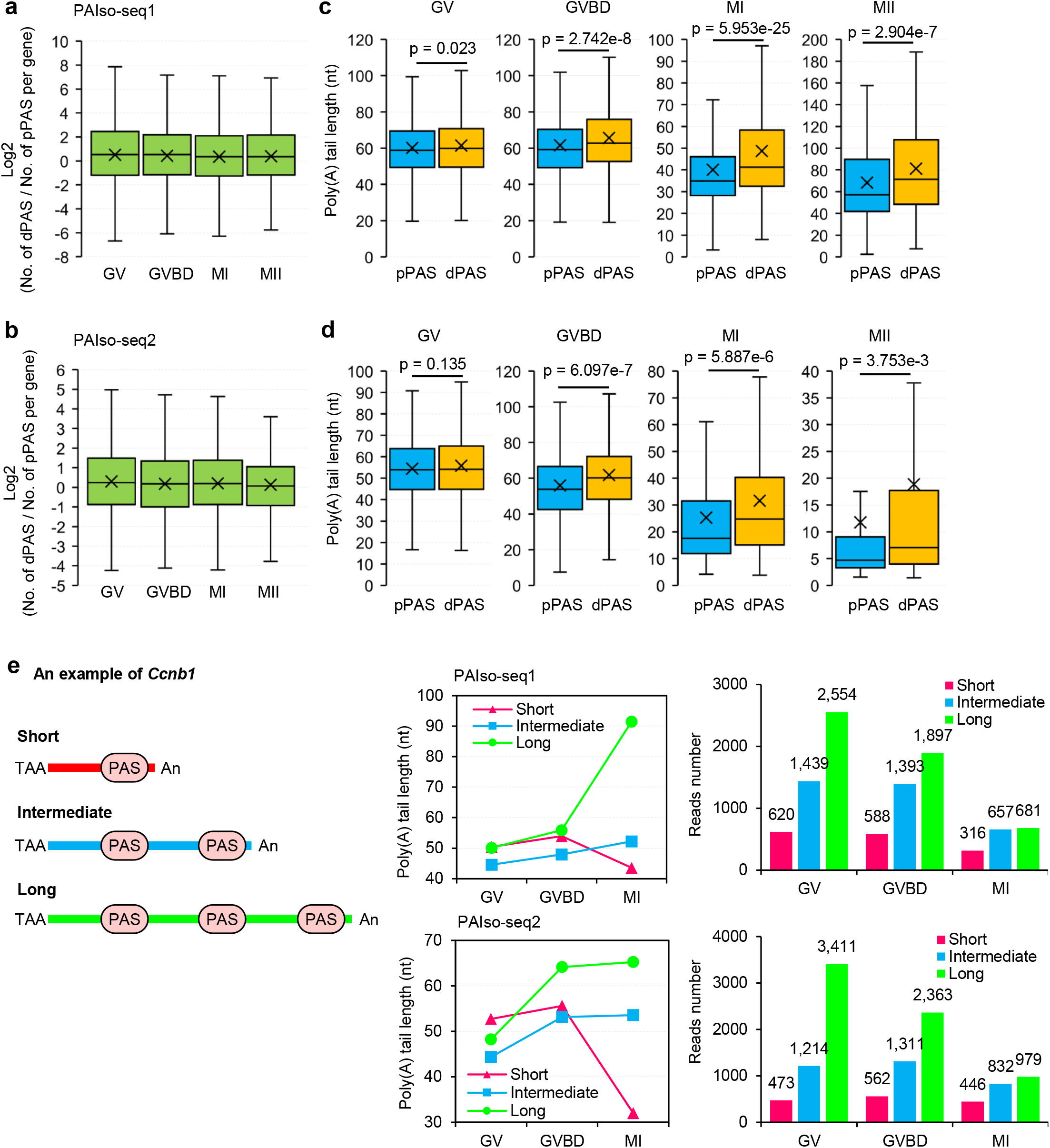
APA isoforms can be differentially regulated at the level of poly(A) tail length in maturing mouse oocytes. **a, b,** Box plots of the ratio of read counts between dPAS (distal PAS) and pPAS (proximal PAS) isoforms per gene on the log2 scale determined from PAIso-seq1 data (**a**) or PAIso-seq2 data (**b**) of mouse oocytes at different stages. Genes with at least 10 poly(A) tail-containing reads (tail length ≥ 1) for both dPAS and pPAS transcripts are included in the analysis. Gene numbers for PAIso-seq1 data (**a**): GV: 1,993, GVBD: 1,432, MI: 1,317, and MII: 488. Gene numbers for PAIso-seq2 data (**b**): GV: 762, GVBD: 591, MI: 494, and MII: 193. **c, d,** Box plots of the poly(A) tail length per isoform for pPAS and dPAS isoforms in PAIso-seq1 data (**c**) or PAIso-seq2 data (**d**) of mouse oocytes at different stages. Genes with at least 10 poly(A) tail-containing reads (tail length ≥ 1) for both dPAS and pPAS transcripts are included in the analysis. *P* value is calculated by Student’s *t*-test. The gene numbers are the same as in (**a**) and (**b**). **e,** Different poly(A) tail dynamics of *Ccnb1* APA isoforms during mouse oocyte maturation. Diagram of *Ccnb1* APA isoforms (left). TAA indicates stop codon. An represents poly(A) tail. PAS indicates polyadenylation signal. Poly(A) tail length dynamics (middle) and read counts (right) of *Ccnb1* isoforms measured by PAIso-seq1 (top) or PAIso-seq2 (bottom). The poly(A) tail length for each PAS isoform is the geometric mean length of all the transcripts with poly(A) tails of at least 1 nt for the given PAS. For all the box plots, the “×” indicates mean value, the black horizontal bars show the median value, and the top and bottom of the box represent the 25^th^ and 75^th^ percentiles, respectively.

Three APA isoforms have been reported for *Ccnb1* in mouse oocytes^45^. The translation of these three isoforms has been shown to be differentially regulated through changes in poly(A) tail length during oocyte maturation^45^. In our mouse PAIso-seq1 and PAIso-seq2 data, good read coverage was obtained for all three isoforms of *Ccnb1* (Fig. 4e). Consistent with the reported trend of change in poly(A) tail length, the shortest isoform of *Ccnb1* showed a gradual decrease in poly(A) tail length, whereas the longest isoform showed an increase in poly(A) tail length; the intermediate isoform also showed an increase, although smaller, in poly(A) tail length during oocyte maturation (Fig. 4e). Given the good correlation between poly(A) tail length and TE in mouse oocytes, regulation of differential TE of APA mRNA isoforms through regulation of poly(A) tail length is likely a general mechanism controlling mammalian oocyte maturation.

### *Btg4* controls active deadenylation during mouse oocyte maturation

Depletion of *Btg4*, which encodes the adaptor of the CCR4-NOT poly(A) tail deadenylase complex, leads to failed OET with developmental arrest between the 1C to 2C stage due to the accumulation of poly(A)+ maternal mRNA^28,30^. The accumulation of maternal mRNA in *Btg4* knockout (KO) oocytes has been linked to impaired mRNA deadenylation of several genes^28,30^. A recent study revealed that BTG4 is also critical for maternal mRNA clearance in human zygotes. *BTG4* mutation in humans leads to accumulation of maternal mRNA in zygotes and developmental arrest of zygotes^46^. However, there is no transcriptome-wide poly(A) tail change evidence for the role of *Btg4* in controlling global maternal mRNA decay through deadenylation. Therefore, we analyzed an *siBtg4* knockdown (KD) MII sample (*siBtg4*) together with a non-targeting siRNA control KD sample (*siNC*) in mouse^47^. Similar to a previous RNA-seq analysis of *Btg4* KO oocytes^30^, we observed accumulation of maternal mRNA in the *Btg4* KD mouse oocytes (Fig. 5a, b). Our data provided us the opportunity to look into the changes in poly(A) tail status after depletion of *Btg4*.

**Fig. 5.**
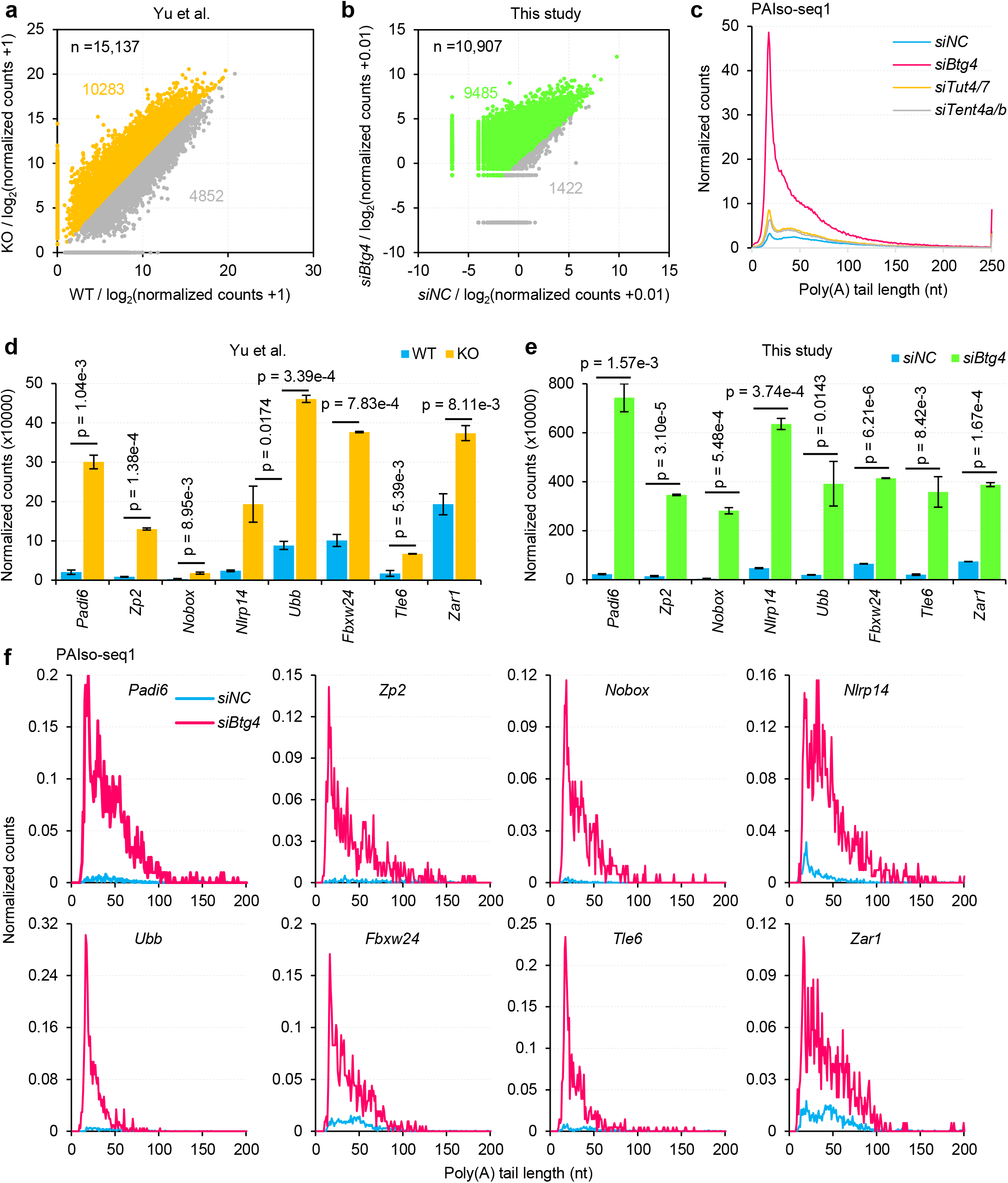
*Btg4* controls maternal mRNA degradation through deadenylation in mouse oocytes. **a,** Scatter plot of the transcriptional levels of individual genes in *Btg4* KO and control MII mouse oocytes from a published RNA-seq dataset^30^. The counts are normalized by reads mapped to protein-coding genes in the mitochondria genome. Each dot represents one gene. The dots in yellow indicate genes with higher normalized counts, while the dots in grey indicate genes with lower normalized counts in *Btg4* KO MII oocytes. **b,** Scatter plot of the transcriptional levels of individual genes in *siBtg4* and control MII mouse oocytes acquired by PAIso-seq1. The counts are normalized by counts of reads mapped to protein-coding genes in the mitochondria genome. Each dot represents one gene. The dots in green indicate genes with higher normalized counts, while the dots in grey indicate genes with lower normalized counts in *Btg4* KD MII oocytes. **c,** Histogram of poly(A) tail lengths of all transcripts in *siNC*, si*Btg4*, *siTut4/7*, or *siTent4a/b* KD mouse oocytes. Histograms (bin size = 1 nt) are normalized by counts of reads mapped to protein-coding genes in the mitochondria genome. Transcripts with a poly(A) tail of at least 1 nt are included in the analysis. Transcripts with poly(A) tail length greater than 250 nt are included in the 250 nt bin. **d, e,** Similar changes in transcriptional abundance of representative genes in *Btg4* KO by RNA-seq (**d**) or *Btg4* KD by PAIso-seq1 (**e**) mouse oocyte samples. Error bars indicate the standard error of the mean (SEM) from two replicates. **f,** Histogram of poly(A) tail lengths of transcripts of the above representative genes (**d** and **e**) in control (*siNC*) and *Btg4* KD (*siBtg4*) MII mouse oocyte samples. Histograms (bin size = 1 nt) are normalized by counts of reads mapped to protein-coding genes in the mitochondria genome. Transcripts with a poly(A) tail of at least 1 nt are included in the analysis. Transcripts with a poly(A) tail length of 1–200 nt are included in the analysis.

Interestingly, global accumulation of polyadenylated transcripts, especially those with short tails, was observed in the *Btg4* KD samples, but not in oocyte with knock-down of the non-canonical poly(A) polymerase *Tut4/7* or *Tent4a/b* (Fig. 5c). For, example, we could see accumulation of *Padi6*, *Zp2*, *Nobox*, *Nlrp14*, *Ubb*, *Fbxw24*, *Tle6*, and *Zar1* mRNA in *Btg4* KD oocytes similar to that in *Btg4* KO oocytes (Fig. 5d, e). These genes showed obvious accumulation of polyadenylated transcripts (Fig. 5f). Given that there is no new transcription during oocyte maturation, the accumulation of polyadenylated transcripts we observed here in *Btg4* KD mouse oocytes must result from defective deadenylation-mediated mRNA decay. Consistent with this, accumulation of polyadenylated transcripts was also observed in our PAIso-seq1 data of the human *BTG4* KD zygotes^48^.

mRNA deadenylation can lead to two outcomes: 1) full deadenylation and decay of transcripts leading to a reduced transcript level and 2) generation of transcripts with relatively short poly(A) tails with the transcript level maintained. Here, we observed high accumulation of transcripts with relatively short poly(A) tails after *Btg4* depletion in mice, but we did not observe accumulation of transcripts with long poly(A) tails. Therefore, it is likely that during mouse oocyte maturation some other factors may first deadenylate mRNA poly(A) tails to a certain length and then the tails are further deadenylated by a Btg4-associated complex, which mainly leads to full deadenylation and decay.

### Conserved extensive mRNA tail re-polyadenylation after fertilization in mice, rats, pigs, and humans

Bulk re-polyadenylation of maternal mRNA in fertilized 1C mouse embryos was initially reported in 1982 based on incorporation of radiolabeled adenosine into existing RNA molecules^49^. Re-polyadenylation of maternal mRNA and new protein translation are essential for zygotic transcriptional competence in mice^50^. However, the molecular identities of the mRNAs that are re-polyadenylated after fertilization during this process are still largely unknown. Our PAIso-seq2 data for mouse MII stage oocytes revealed a large number of mRNAs without poly(A) tails or with very short poly(A) tails, which we hypothesized to be potential substrates for maternal mRNA recruitment and re-polyadenylation following fertilization^50^. To address this on a transcriptome-wide scale, we compared the transcriptome-wide poly(A) tails before and after fertilization using both the PAIso-seq1 and PAIso-seq2 methods (Fig. 6a). PAIso-seq1 uses templated end-extension to add adaptors to the mRNA 3’-ends, which requires a stretch of A residues at the mRNA 3’-end; therefore, mRNAs without poly(A) tails cannot be sequenced by PAIso-seq1, and mRNAs with short poly(A) tails are sequenced less efficiently (Fig. 6b). PAIso-seq2 employs ligation of an adaptor to the mRNA 3’-ends, which is not affected by the length or composition of the mRNA poly(A) tails (Fig. 6b). Given that there is no new transcription in MII oocytes and 1C embryos immediately after fertilization, we can take advantage of the features of PAIso-seq1 and PAIso-seq2 to analyze the global polyadenylation status of mRNA before and after fertilization based on the following reasoning: PAIso-seq1 detection relies on poly(A) tails, and can be used quantify the relative amount of transcripts with poly(A) tails, while PAIso-seq2 detection is independent of poly(A) tails and can serve as a control for the total transcript level and be used for quantification of transcripts without poly(A) tails.

**Fig. 6.**
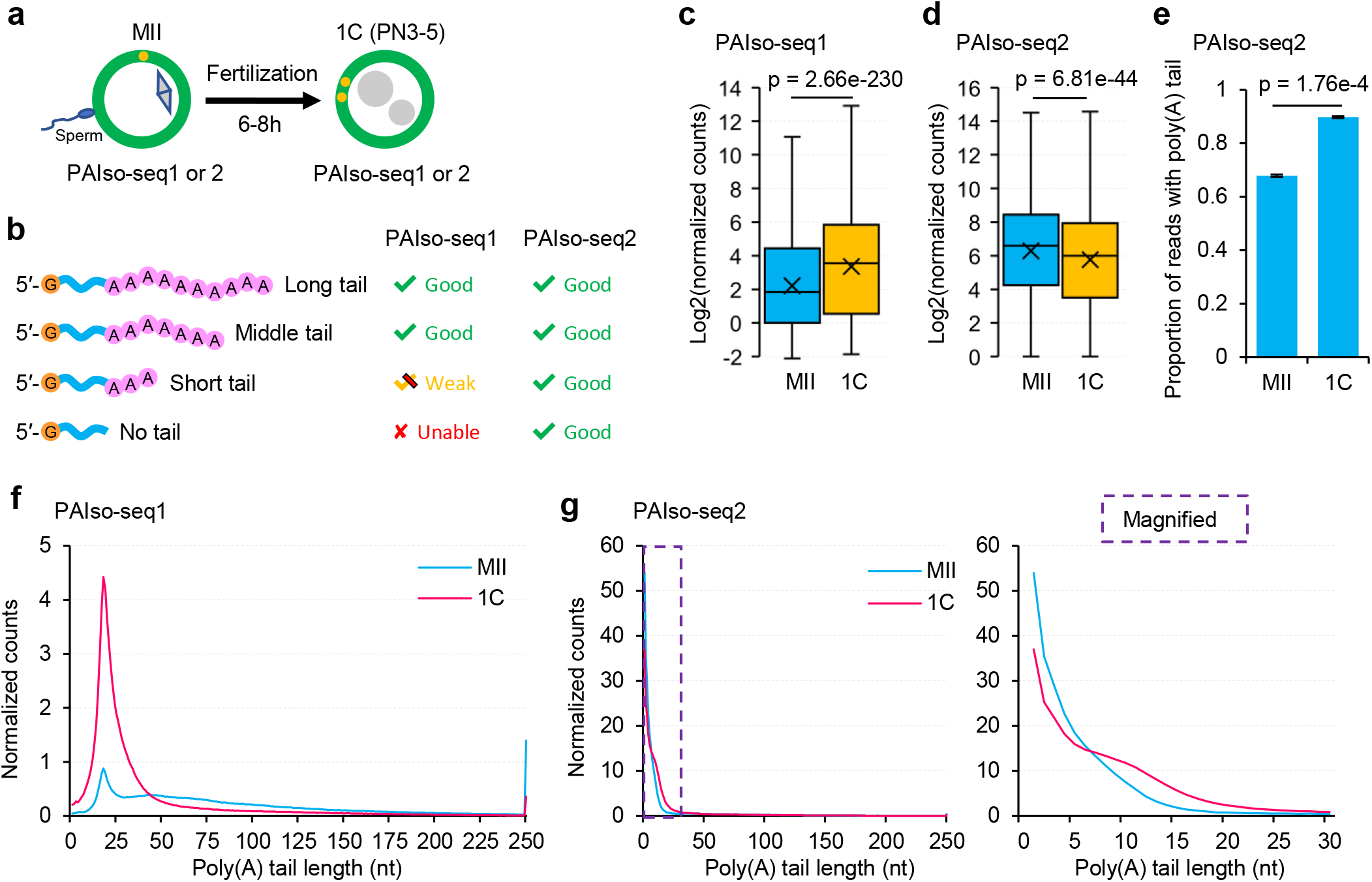
Extensive mRNA tail re-polyadenylation after fertilization in mouse. **a,** Diagram of experiments performed to analyze poly(A) tail re-polyadenylation after fertilization. **b,** Diagram showing the different abilities of PAIso-seq1 and PAIso-seq2 to sequence poly(A) tails of different length. **c, d,** Box plots of the normalized gene expression levels of individual genes in mouse MII oocyte and 1C embryo samples sequenced by PAIso-seq1 (**c**) or PAIso-seq2 (**d**). The read counts for individual genes are normalized by counts of reads mapped to protein-coding genes in the mitochondria genome. *P* value is calculated by Student’s *t*-test. The “×” indicates mean value, the black horizontal bars show the median value, and the top and bottom of the box represent the 25^th^ and 75^th^ percentiles, respectively. **e,** Proportion of sequenced transcripts containing poly(A) tails in mouse MII oocytes and 1C embryos, as determined by PAIso-seq2. Transcripts with a poly(A) tail of at least 1 nt are considered reads with poly(A) tails. Error bars indicate the SEM from two replicates. *P* value is calculated by Student’s *t*-test. **f,** Histogram of poly(A) tail lengths of all transcripts in mouse MII oocytes and 1C embryos sequenced by PAIso-seq1. Histograms (bin size = 1 nt) are normalized by counts of reads mapped to protein-coding genes in the mitochondria genome. Transcripts with a poly(A) tail of at least 1 nt are included in the analysis. Transcripts with a poly(A) tail length greater than 250 nt are included in the 250 nt bin. **g,** Histogram of poly(A) tail lengths of all transcripts in mouse MII oocytes and 1C embryos sequenced by PAIso-seq2. Magnified view (region in the purple dotted square) of short poly(A) tails is shown on the right. Histograms (bin size = 1 nt) are normalized by counts of reads mapped to protein-coding genes in the mitochondria genome. Transcripts with a poly(A) tail of at least 1 nt are included in the analysis. Transcripts with a poly(A) tail length greater than 250 nt are included in the 250 nt bin.

We observed that in mouse the level of transcripts with poly(A) tails for each gene detected by PAIso-seq1 significantly increased in 1C embryos compared with MII oocytes (Fig. 6c), while the total level of the transcripts did not increase (Fig. 6d). Consistent with the above results, the fraction of transcripts with poly(A) tails significantly increased after fertilization of mouse oocytes as measured by PAIso-seq2 (Fig. 6e). Looking at the distribution of mRNA poly(A) tails, we observed the appearance of a sharp peak of short poly(A) tails after fertilization as measured by PAIso-seq1 data (Fig. 6f). The poly(A) tail length distribution of 1C embryos after fertilization shifted toward longer poly(A) tails compared with that of MII oocytes as determined using PAIso-seq2 data (Fig. 6g). The PAIso-seq1 and PAIso-seq2 datasets are highly consistent. The mRNA poly(A) tails are re-polyadenylated resulting in longer tails, which allows them to be sequenced by PAIso-seq1, resulting in the sharp peak of short poly(A) tails in the mouse PAIso-seq1 data.

Next, we asked whether the global mRNA re-polyadenylation after fertilization is specific to mice or conserved in mammals. To address this, we analyzed our transcriptome-wide poly(A) tail data for rats, pigs, and humans. In rats, there were minimal changes in the transcript levels of each gene after fertilization as measured by PAIso-seq2 (Fig. 7a). Similar to what we observed in mouse, we observed a significantly increased proportion of polyadenylated transcripts (Fig. 7b), as well as a shift in the global poly(A) tail length distribution toward longer tails (Fig. 7c). In pigs, we observed an increased level of transcripts for each gene using PAIso-seq1 but not PAIso-seq2 (Fig. 7d, e). We also observed an increased proportion of polyadenylated transcripts together with increased global poly(A) tail length, which is almost identical to the pattern seen in mouse (Fig. 7f-h). In humans, we observed a similar trend for transcript level of each gene: an increase in the PAIso-seq1 data but not in the PAIso-seq2 data (Fig. 7i, j). An increased proportion of polyadenylated transcripts was also seen in human embryos after fertilization, although this increase was delayed until the 2C stage (Fig. 7k). Similarly, we observed a global increase in poly(A) tail length after fertilization in humans (Fig. 7l, m).

**Fig. 7.**
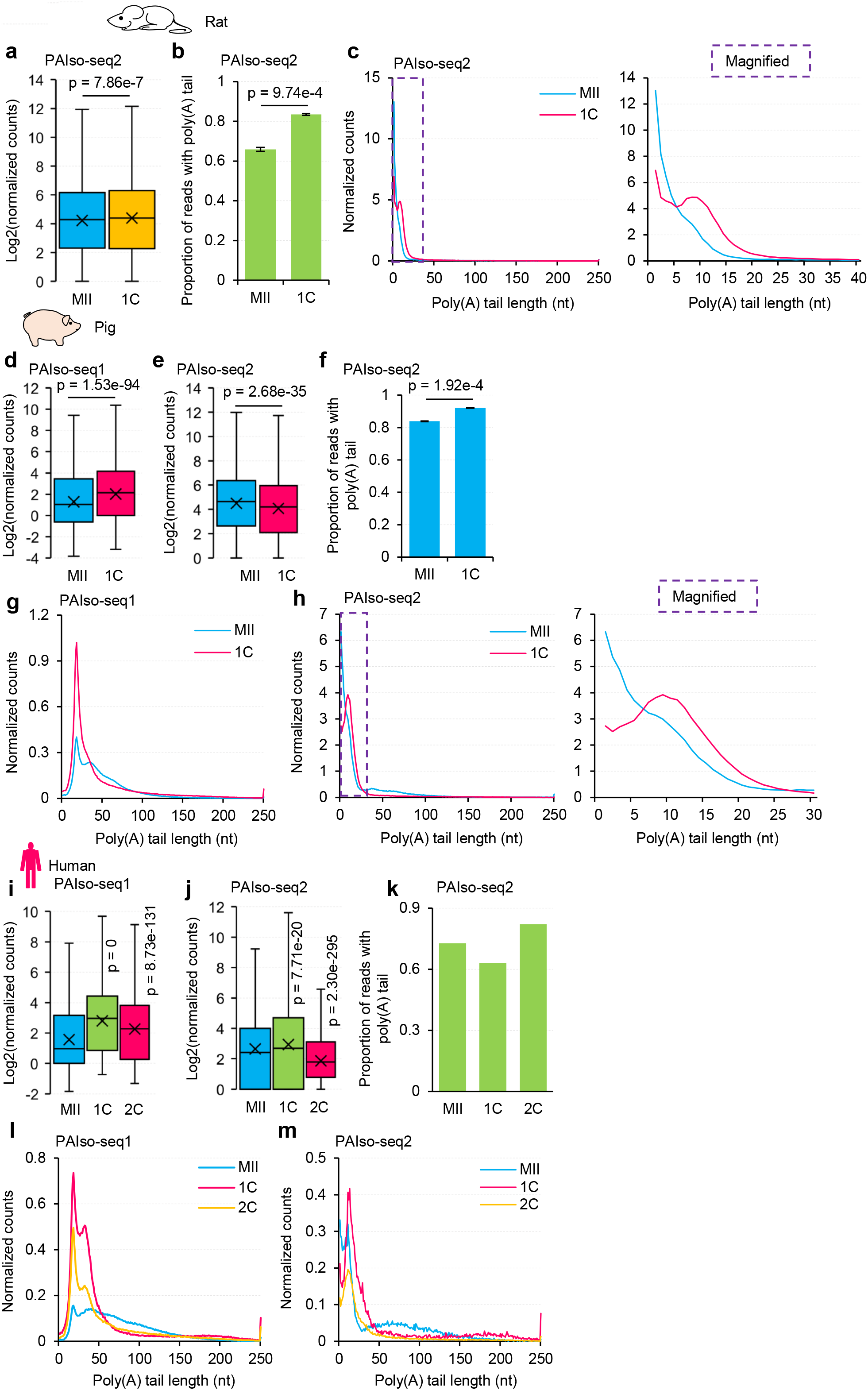
Global RNA re-polyadenylation after fertilization is conserved in rats, pigs, and humans. **a,** Box plot of the normalized gene expression levels of individual genes in rat MII oocyte and 1C embryo samples sequenced by PAIso-seq2. **b,** Proportion of transcripts sequenced by PAIso-seq2 containing poly(A) tails in rat MII oocytes and 1C embryos. Transcripts with a poly(A) tail of at least 1 nt are considered reads with poly(A) tails. Error bars indicate the SEM from two replicates. **c,** Histogram of poly(A) tail lengths of all transcripts in rat MII oocytes and 1C embryos sequenced by PAIso-seq2. Magnified view (region in the purple dotted square) of short poly(A) tails is shown on the right. **d, e,** Box plot of the normalized gene expression level of individual genes in pig MII oocyte and 1C embryo samples sequenced by PAIso-seq1 (**d**) or PAIso-seq2 (**e**). **f,** Proportion of transcripts sequenced by PAIso-seq2 containing poly(A) tails in pig MII oocytes and 1C embryos. Transcripts with a poly(A) tail of at least 1 nt are considered reads with poly(A) tails. Error bars indicate the SEM from two replicates. **g,** Histogram of poly(A) tail lengths of all transcripts in pig MII oocytes and 1C embryos sequenced by PAIso-seq1. **h,** Histogram of poly(A) tail lengths of all transcripts in pig MII oocytes and 1C embryos sequenced by PAIso-seq2. Magnified view (region in the purple dotted square) of short poly(A) tails is shown on the right. **i, j,** Box plot of the normalized gene expression levels of individual genes in human MII oocyte and 1C and 2C embryo samples sequenced by PAIso-seq1 (**i**) or PAIso-seq2 (**j**). **k,** Proportion of transcripts sequenced by PAIso-seq2 containing poly(A) tails in human MII oocyte and 1C and 2C embryo samples. Transcripts with a poly(A) tail of at least 1 nt are considered reads with poly(A) tails. **l, m,** Histogram of poly(A) tail lengths of all transcripts in human MII oocyte and 1C and 2C embryo samples sequenced by PAIso-seq1 (**l**) or PAIso-seq2 (**m**). The read counts are normalized by counts of reads mapped to protein-coding genes in the mitochondria genome if normalization is indicated. For all the box plots, the “×” indicates mean value, the black horizontal bars show the median value, and the top and bottom of the box represent the 25^th^ and 75^th^ percentiles, respectively. *P* value is calculated by Student’s *t*-test. Histograms (bin size = 1 nt) include transcripts with a poly(A) tail of at least 1 nt. For histograms with an X axis upper limit of 250, transcripts with poly(A) tail lengths greater than 250 nt are included in the 250 nt bin.

Taken together, our data reveal the conserved transcriptome-wide global re-polyadenylation of mRNA poly(A) tails after fertilization in mice, rats, pigs, and humans, resulting in accumulation of re-polyadenylated transcripts with short poly(A) tails. The re-polyadenylated mRNA can be protected by G residues incorporated into the poly(A) tails by Tent4a/b in both mouse oocytes and human zygotes^47,48^. In addition, the fertility was reduced in *Tent4a* KO female mice (our unpublished observation). Together with the findings in the current study, these suggest that maternal mRNA regulation through re-polyadenylation is essential for zygote development.

## Discussion

RNA poly(A) tails were discovered in 1971, and are now understood to be one of the most important functional components of mature mRNA^51–53^. Transcription is silent during the mammalian OET before ZGA, and regulation via mRNA poly(A) tails is thus a major post-transcriptional regulation mechanism ensuring proper regulation of diverse events during the OET. In addition, mammalian oocytes provide a straightforward experimental system for studying mRNA poly(A) tail regulation without complications arising from new transcription. In the present study, we revealed the dynamic landscape of mRNA poly(A) tails throughout the mouse OET using two methods based on different underlying principles. We found that global mRNA poly(A) tail length dynamics shape the translational profiles during mouse oocyte maturation. A large fraction of maternal mRNAs follows a general trend of shortening of poly(A) tails during oocyte maturation (Fig. 8a). In addition, we revealed that Btg4-mediated deadenylation serves as an important pathway for maternal mRNA decay, which is essential for oocyte maturation and early embryo development (Fig. 8a). Finally, we revealed conserved transcriptome-wide re-polyadenylation after fertilization in mice, rats, pigs, and humans, which reshapes the transcriptome for pre-implantation embryo development (Fig. 8a).

**Fig. 8.**
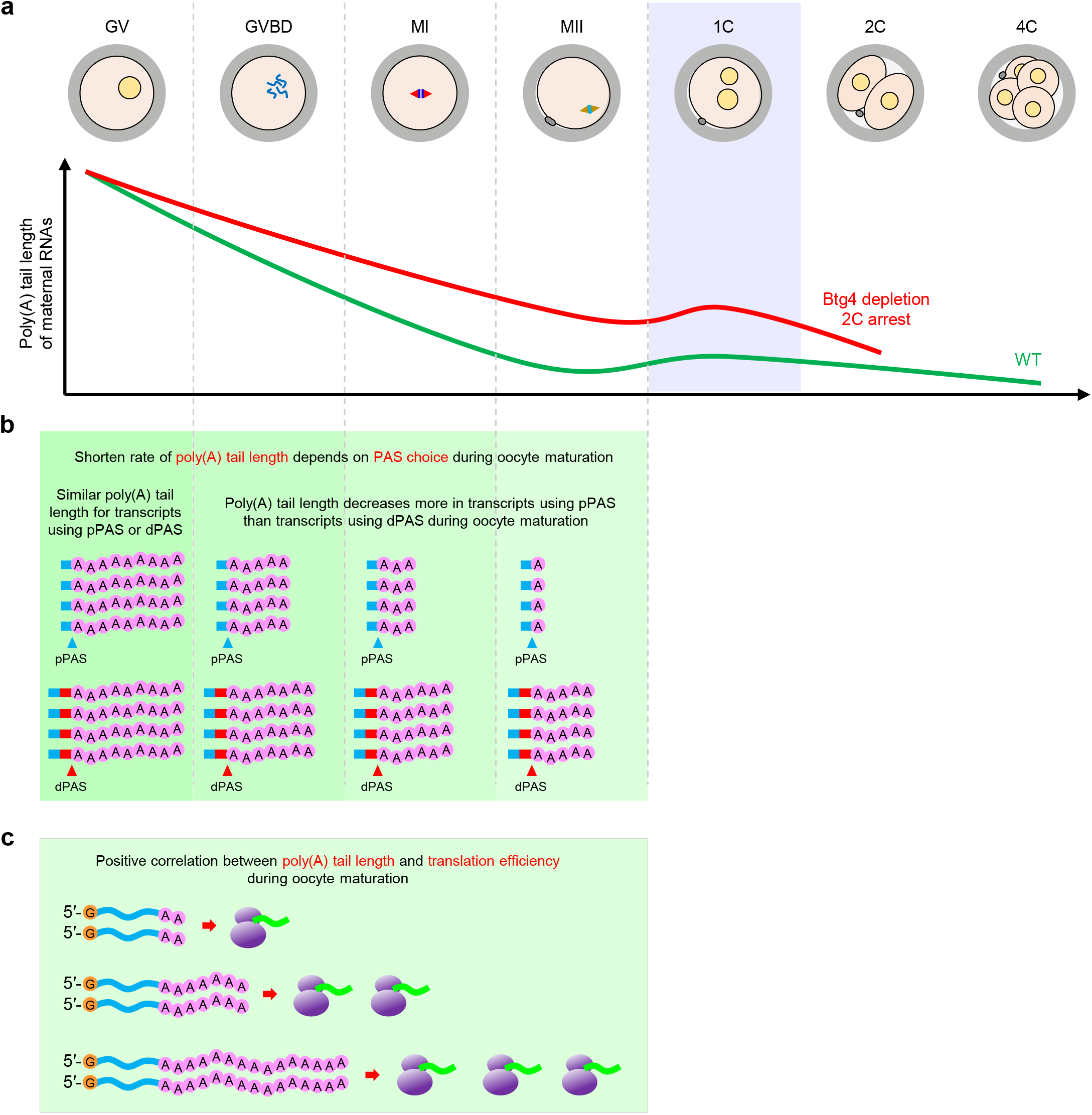
A model of poly(A) tail length dynamics and its association with translation regulation during mammalian OET. **a,** A model of poly(A) tail length dynamics (green line) during the mouse oocyte-to-embryo-transition. One additional line (red) shows poly(A) tail length dynamics in *Btg4*-depleted oocytes and embryos. The global mRNA re-polyadenylation event after fertilization is highlighted in violet. **b,** A model of differential regulation of poly(A) tails in transcripts with proximal and distal PASs during oocyte maturation. In general, poly(A) tails of pPAS transcripts undergo more extensive deadenylation than those of dPAS transcripts. **c,** A model of the positive association between poly(A) tail length and translation, in which mRNAs with longer poly(A) tails are translated more efficiently.

We noticed that in maturing mouse, rat, pig, and human oocytes, the dPAS isoforms had longer poly(A) tails compared with pPAS isoforms. Given that mRNA poly(A) tail length associates well with translational efficiency in mouse oocytes, it is likely that these dPAS isoforms are preferentially translated in maturing mammalian oocytes. In contrast, this phenomenon is not seen in 3T3 or mouse embryonic stem cells^54^. Therefore, this phenomenon represents a novel general regulatory mechanism for isoforms generated through alternative polyadenylation in maturing oocytes. As there is no new transcription during oocyte maturation, the observed difference in poly(A) tail dynamics between the pPAS and dPAS isoforms must result from post-transcriptional regulation through poly(A) tails. The poly(A) tails of pPAS and dPAS isoforms are of similar length in the GV stage. We noticed that the pPAS isoforms follow a deadenylation trend similar to the global deadenylation trend during oocyte maturation, whereas the deadenylation of dPAS isoforms is much slower than that of pPAS isoforms (Fig. 8b, c). It is likely that cis-elements present in the longer 3’-UTRs of dPAS isoforms may account for the regulatory specificity, and this warrants further investigation in the future.

We observed global re-polyadenylation in mouse, rat, pig, and human zygotes after fertilization. When this re-polyadenylation was blocked by treatment with 3’-dA (cordycepin) (Extended data Fig. 3a), an adenosine analog that can be incorporated into RNA poly(A) tails and prevent further elongation due to the absence of a 3’ hydroxyl group^50,55^, all of the treated mouse embryos were arrested at the 2C stage (Extended data Fig. 3b, c). Similarly, treating human zygotes with 3’-dA immediately after intracytoplasmic sperm injection resulted in zygotic cleavage failure (Extended data Fig. 3i). At the molecular level, the poly(A) tail length and the polyadenylated transcript level detected by PAIso-seq1 decreased after 3’-dA treatment in both mouse and human embryos (Extended data Fig. 3d-g, j, k). In addition, fertility was reduced in female mice depleted of *Tent4a* (our unpublished observation), a non-canonical poly(A) polymerase, further supporting the importance of poly(A) tail-mediated regulation of zygote development. It has been shown that re-polyadenylation of maternal mRNA and subsequent new protein translation are essential for mouse zygotic transcriptional competence^50^. We revealed the appearance of a large number of mRNAs with short poly(A) tails in mouse, rat, pig, and human zygotes. These tails may be sufficient to support translation of the corresponding mRNAs as shown in a reporter assay (Extended Data Fig. 3h). Therefore, mRNA re-polyadenylation after fertilization is critical for embryo development in mammals. Elucidation of the detailed functions of the re-polyadenylated mRNAs is an interesting direction for exploration in the future.

In flies, global cytoplasmic re-polyadenylation during late oogenesis and early embryogenesis is essential for the egg-to-embryo transition, and requires Wispy, a Gld-2 type non-canonical poly(A) polymerase^17,18,56,57^. Tent2/GLD2 in vertebrates can also catalyze cytoplasmic polyadenylation^58,59^. It was therefore hypothesized that Tent2 in mice could catalyze global re-polyadenylation following fertilization, similar to Wispy in flies. However, a *Tent2* maternal KO displayed normal fertility without detectable changes in RNA polyadenylation activity in MII-stage oocytes, which is evidence against Tent2 involvement in global re-polyadenylation after fertilization^60^. This raises the interesting question of which enzyme contributes to the global re-polyadenylation after fertilization in mammals. We showed that Tent4a/b can re-polyadenylate and stabilize poly(A) tails through incorporation of G modifications in mouse oocytes and human zygotes^47,48^. Therefore, which enzyme is responsible for the re-polyadenylation after fertilization, or which enzyme re-polyadenylates maternal mRNA together with Tent4a, is a very interesting question to be answered in the future.

In summary, our study reveals the previously unseen highly conserved transcriptome-wide dynamics of mRNA poly(A) tail length during OET in mice, rats, pigs, and humans, offering an unprecedented high-resolution data resource that opens the door for a new era of post-transcriptional RNA regulatory studies of the mammalian OET.

## Materials and Methods

### Animals and collection of embryos

Mice were purchased from Beijing Vital River Laboratory Animal Technology Co., Ltd., and used in compliance with the guidelines of the Animal Care and Use Committee of the Institute of Genetics and Development Biology, Chinese Academy of Sciences. For preparation of 1C embryos, 7–8-week-old CD1 (ICR) mice were superovulated by intraperitoneally injecting them with 10 U of pregnant mare serum gonadotropin (PMSG, Prospec). Fertilized eggs surrounded by cumulus cells were collected from the oviduct 12 hours after human chorionic gonadotrophin (hCG, Prospec) injection. The pronuclei had yet not formed at this moment. Some unfertilized eggs were also recovered and were later discarded. The collected embryos were ready for the 3’-dA treatment and other experiments. **Drug treatment of oocytes and embryos**

The drug 3’-dA (Sigma) was dissolved in M16 medium (Sigma). Medium without the drug was used as a control. For treatment of 1C embryos, embryos were cultured directly in medium with or without 3’-dA. For treatment of mouse embryos, 3’-dA was added to the embryo culture medium prior to the formation of the pronucleus. For the treatment of human zygotes, zygotes were cultured directly in medium containing 3’-dA immediately after intracytoplasmic sperm injection.

### Microinjection of mRNA

The templates used for *in vitro* transcription were amplified by PCR from the pcDNA3.1-EGFP/mCherry-3XFlag plasmids using the T7-F forward primer and different reverse primers (Extended Data Table S1). The capped mRNAs were *in vitro* synthesized with the HiScribe T7 ARCA mRNA Kit (NEB) and purified with an RNA Clean & Concentrator-25 kit (Zymo Research) according to the manufacturers’ instructions. The purified mRNA was dissolved in RNase-free water and stored at -80 ℃ until use. The GV oocytes were microinjected with 5–10 pl mRNA (200 ng/μl) and cultured for the indicated time.

### Total RNA isolation

For the mouse embryos, total RNA was extracted with Direct-zol RNA MicroPrep (Zymo Research) according to the manufacturer’s instruction. Briefly, embryo were lysed directly in 500 μl or 1ml of TRIzol reagent (Ambion) and mix thoroughly, then 500 μl or 1 ml of 100% ethanol was added and mixed thoroughly. The mixture was transferred into a Zymo-Spin IC Column and centrifuged to capture the RNA on the column. After washing, the RNA was eluted by adding 15-40 μl RNase-free water directly to the column matrix. Repeat the elution. Prepared RNA was stored at -80℃ or used immediately.

### PAIso-seq1 and PAIso-seq2 library construction

Total RNA from mouse embryos were used for PAIso-seq1 as described previously^4^. For human embryos, PAIso-seq2 libraries were constructed with 4 or 5 oocytes/embryos following the PAIso-seq2 protocol described in another study^5,48^. The libraries were size selected by Pure PB beads (1x beads for cDNA more than 200 bp and 0.4x beads for cDNA more than 2 kb, the two parts of the sample were combined at equal molar for further library construction), made into SMRTbell Template libraries (SMRTbell Template Prep Kit). The libraries were annealed with the sequencing primer and bind to polymerase, finally the polymerase-bound template was bound to Magbeads and sequenced using PacBio Sequel or Sequel II instruments at Annoroad.

### PAIso-seq1 sequencing data processing

The PAIso-seq1 data generated in this study was processed following the methods described^47^. In brief, to demultiplex and extract the transcript sequence from the CCS reads, we firstly matched the barcodes in the CCS reads and the reverse complementary of the CCS reads allowing maximum two mismatches or indels (insertions and deletions). CCS reads were oriented and split into multiple transcripts if multiple barcodes were matched. Then, to get the precise 3’ end position of the original RNA, we aligned the matched barcode to each transcript using a semi-global function “sg_dx_trace” which did not penalize gaps at both the beginning and the end of query/barcode in parasail package^61^ and trimmed the barcode. Finally, the 3’-adapter and the 5’-adapter of each transcript were removed. Transcripts with length greater than 50nt were kept. Clean CCS reads were used for the downstream analysis.

Clean CCS reads were aligned to the reference genome using minimap2 v.217-r941^62^ with parameters “-ax splice -uf --secondary=no -t 40 -L --MD --cs --junc-bed mm10.junction.bed”. The mm10.junction.bed file was converted from GRCm38 mouse gene annotation with “paftools.js gff2bed” in the minimap2 package. Read counts of each gene and gene assignments of each CCS reads were summarized by featureCounts v2.0.0^63^ with “-L -g gene_id -t exon -s 1 -R CORE -a Mus_musculus.GRCm38.92.gtf” parameters using the read alignments generated by minimap2. Now the clean CCS reads are ready for downstream analysis.

### PAIso-seq2 data pre-processing

The PAIso-seq2 data generated in this study was processed following the methods described^5^. In brief, to demultiplex and extract the transcript sequence from the CCS reads, we firstly matched the barcodes in the CCS read and the reverse complementary of the CCS reads allowing maximum two mismatches or indels (insertions and deletions). CCS reads were oriented and split into multiple transcripts if multiple barcodes were found. Then, to get the precise 3’ end position of the original RNA, we aligned the matched barcode to each transcript using a semi-global function “sg_dx_trace” which did not penalize gaps at both the beginning and the end of query/barcode in parasail package^61^ and trimmed the barcode. Next, we used the following regular pattern “(AAGCAGTGGTATCAACGCAG){e<=2}(AGTAC){s<=1}([ATCG]{8,12})(ATGGG) {s<=1}” to match the 5’-adapter and extract the UMIs in each transcript. Finally, the 3’-adapter and the 5’-adapter of each transcript were removed and the remaining extracted sequence was called clean CCS hereafter. Clean CCS reads were used for the downstream analysis.

Clean CCS reads alignment to the reference genome and gene assignment were performed the same way as described in the PAIso-seq1 data pre-processing. Clean CCS reads with the identical mapping position (namely, the start and end position mapped to the reference genome) and the identical UMI (Unique Molecular Identifiers) sequence were determined and only one clean CCS reads was kept. Now the clean CCS reads are ready for downstream analysis.

### Poly(A) tail sequence extraction

Clean CCS reads were pre-processed as described above. Minimap2 alignments with “SA” (supplementary alignment) tag were ignored. The terminal clipped sequence of the CCS reads in the alignment bam file is used as candidate poly(A) tail sequence. We defined a continuous score based on the transitions between the two adjacent nucleotide residues throughout the 3’-soft clip sequences. To calculate continuous score, the transition from one residue to the same residue scored 0, and the transition from one residue to a different residue score 1. Number of A, U, C and G residues were also counted in the 3’-soft clip sequences of each alignment. The 3’-soft clip sequences with the frequency of U, C and G greater or equal to 0.1 simultaneously were marked as “HIGH_TCG” tails. The 3’-soft clips which were not marked as “HIGH_TCG” and with continuous score less than or equal to 12 were considered as poly(A) tails.

### Poly(A) tail length measurement

To determine the lengths of poly(A) tail accurately, we only quantify the poly(A) tail length from clean CCS reads with at least ten passes. The poly(A) tail length of a transcript was calculated as the length of the sequence, including U, C or G residues if present. The poly(A) tail length of a gene was presented by the geometric mean of the poly(A) tail length of transcripts with tail length at least 1 nt from the given gene, because poly(A) tail length distribution of a gene is a lognormal-like distribution^18^.

### Analysis of maternal and zygotic genes

The maternal and zygotic genes was defined using the PAIso-seq1 or PAIso-seq2 data following the published strategy with minor modification^64^. In brief, edgeR was used for differential expression analysis^65^. The maternal genes were defined by protein coding genes with showing 4-fold enrichment in GV oocytes compared to 2C-embryos at the transcript level and a p value < 0.05, while the zygotic genes were defined by protein coding genes with showing 4-fold enrichment in 2C embryos compared to GV oocytes at the transcript level and a *p* value < 0.05.

### Genome and gene annotation

The genome sequence used in this study is from the following links. ftp://ftp.ensembl.org/pub/release-92/fasta/mus_musculus/dna/Mus_musculus.GRCm38.dna_rm.primary_assembly.fa.gz. The genome annotation (including the nuclear encoded mRNAs, lncRNAs and mitochondria encoded mRNAs) used in this study is from the following links. ftp://ftp.ensembl.org/pub/release-92/gtf/mus_musculus/Mus_musculus.GRCm38.92.gtf.gz

### Data Availability

The ccs data in bam format from PAIso-seq1 and PAIso-seq2 experiments will be available at Genome Sequence Archive hosted by National Genomic Data Center. Other datasets used in this paper include: Luong et al. (Gene Expression Omnibus database (GEO) no. GSE135525) used in Fig. 2, 3; Yu et.al. (GEO no. GSE71257) used in Fig. 5a, d. Custom scripts used for data analysis will be available upon request.

## Acknowledgements

We thank Hongxiang Liu for his technical assistance in mouse superovulation and collection of oocytes and embryos. We thank Yiwei Zhang for his technical assistance in bioinformatic analysis. This work was supported by the National Key Research and Development Program of China (2018YFA0107001), the Strategic Priority Research Program of the Chinese Academy of Sciences (XDA24020203), National Natural Science Foundation of China (31970588, 32170606), Natural Science Foundation of Heilongjiang province (YQ2020C003), the China Postdoctoral Science Foundation (2020M670516, 2020T130687), the State Key Laboratory of Molecular Developmental Biology, and the Fundamental Research Funds of Shandong University.

## Author Contributions

Yusheng Liu, Jiaqiang Wang and Falong Lu conceived the project and designed the study. Yusheng Liu collected and performed drug treatment on mouse embryos, mRNA preparation and microinjection in mouse oocytes. Chuanxin Zhang and Zhenzhen Hou collected and performed drug treatment on human embryos. Yusheng Liu, Hu Nie, Jiaqiang Wang and Falong Lu analyzed the sequencing data. Yusheng Liu and Jiaqiang Wang organized all figures. Yusheng Liu, Jiaqiang Wang and Falong Lu supervised the project. Yusheng Liu, Jiaqiang Wang and Falong Lu wrote the manuscript with the input from the other authors.

## Competing Interests statement

The authors declare no competing interests.

---

**Fig. S1.**
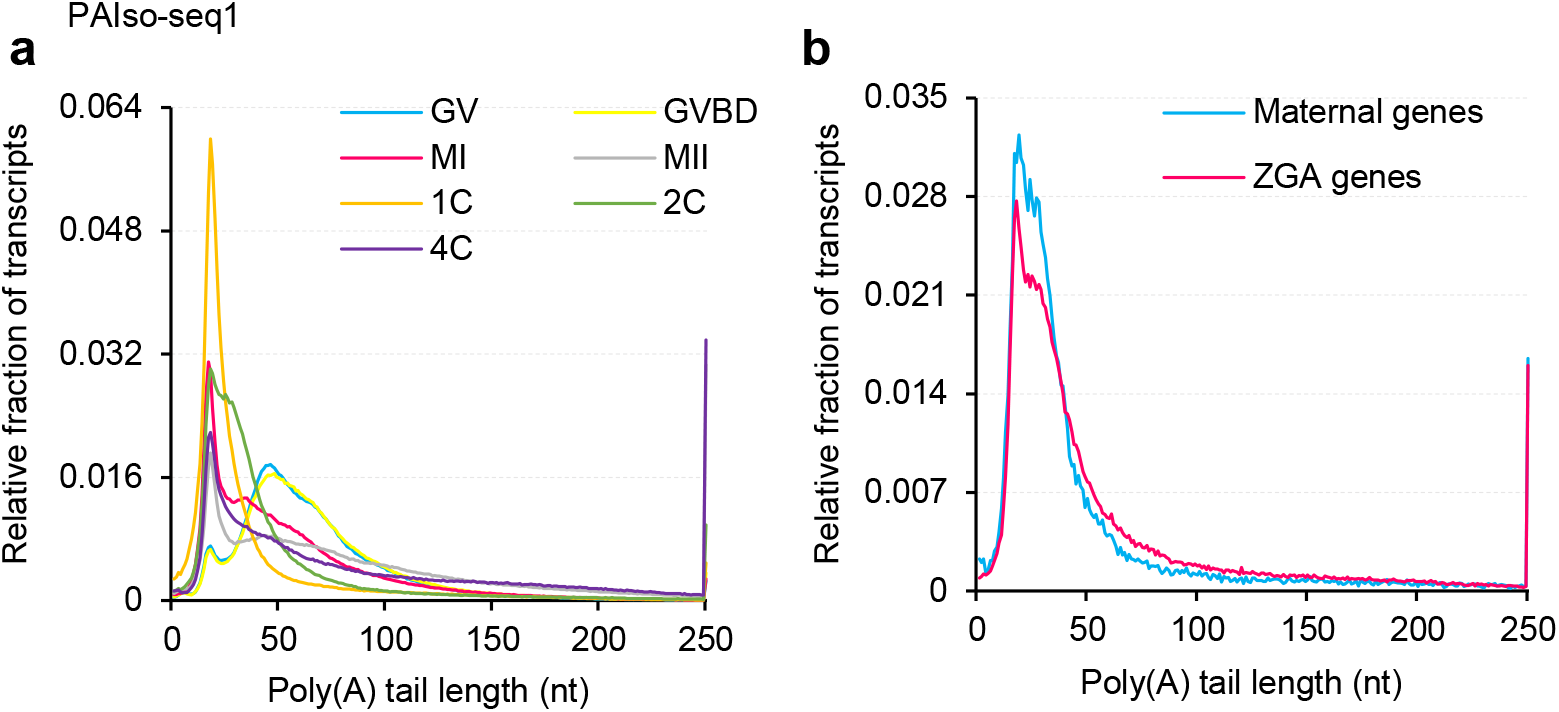
Dynamics of mRNA poly(A) tail length during the mouse OET revealed by PAIso-seq1. **a,** Histogram of transcriptome-wide poly(A) tail lengths in mouse oocytes and early embryos measured by PAIso-seq1. **b,** Histogram of poly(A) tail lengths of combined transcripts from maternal genes and zygotic genes measured by PAIso-seq1 in mouse 2-cell stage embryos. Histograms (bin size = 1 nt) are normalized to cover the same area. Transcripts with a poly(A) tail of at least 1 nt are included in the analysis. Transcripts with poly(A) tail length greater than 250 nt are included in the 250 nt bin.

**Fig. S2.**
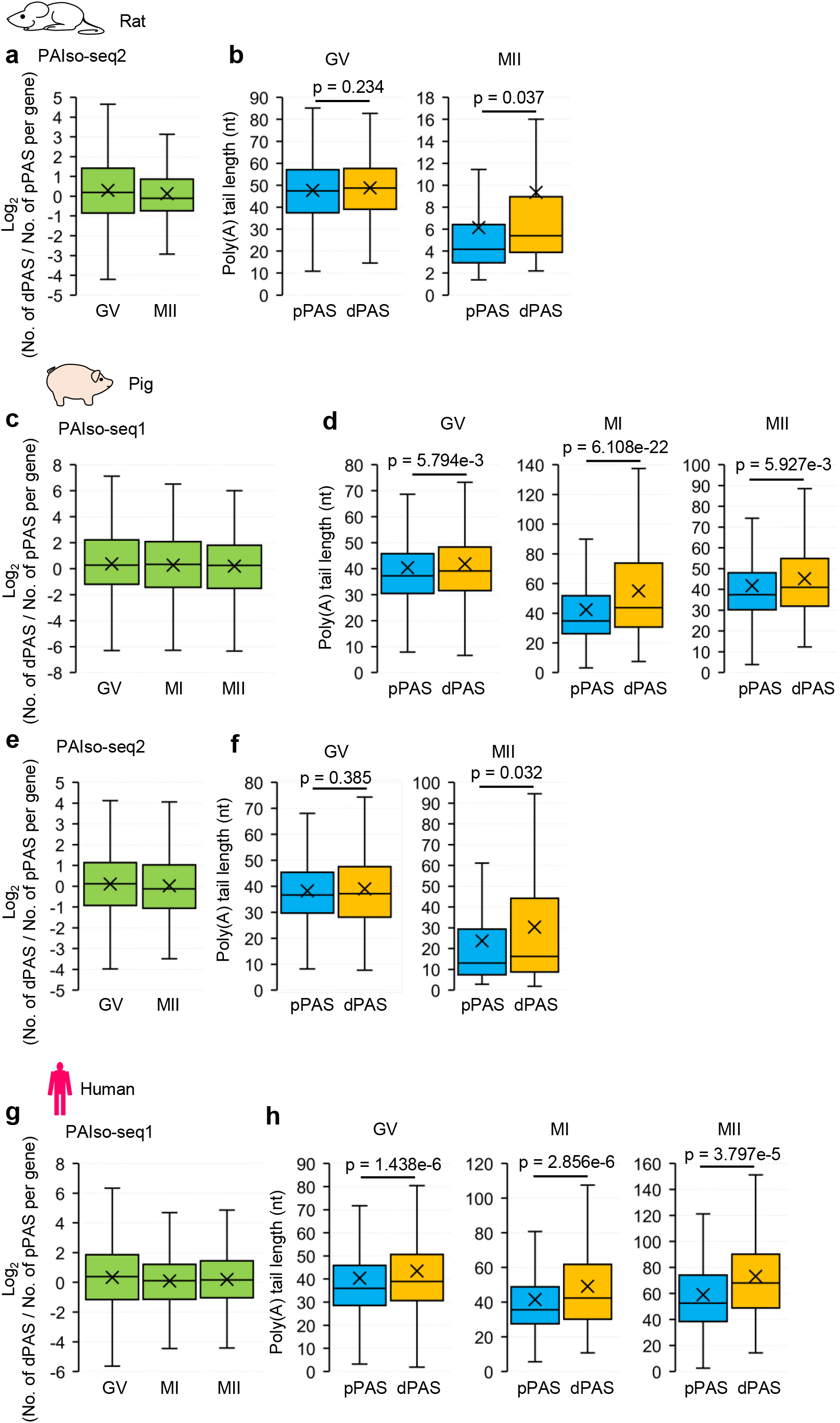
Differential poly(A) tail length dynamics between APA isoforms is conserved among rats, pigs and humans. **a,** Box plot of the ratio of read counts between dPAS (distal PAS) and pPAS (proximal PAS) isoforms per gene on the log2 scale determined from PAIso-seq2 data for rat oocytes at different stages. Genes (GV: *n* = 356, MII: *n* = 50) with at least 10 poly(A) tail-containing reads (tail length ≥ 1) for both dPAS and pPAS isoforms are included in the analysis. **b,** Box plots of the poly(A) tail length per isoform for pPAS and dPAS isoforms determined from PAIso-seq2 data for rat oocytes at different stages. Gene number: GV: 356, MII: 50. **c,** Box plot of the ratio of read counts between dPAS and pPAS isoforms per gene on the log2 scale determined from PAIso-seq1 data from pig oocytes at different stages. Gene number: GV: 1,932, MI: 948, MII: 476. **d,** Box plot of the poly(A) tail length per isoform for pPAS and dPAS isoforms determined from PAIso-seq1 data of pig oocytes at different stages. Gene number: GV: 1,932, MI: 948, MII: 476. **e,** Box plot of the ratio of read counts between dPAS and pPAS isoforms per gene on the log2 scale determined from PAIso-seq2 data of pig oocytes at different stages. Gene number: GV: 562, MII: 174. **f,** Box plot for the poly(A) tail length per isoform for pPAS and dPAS isoforms determined from PAIso-seq2 data of pig oocytes at different stages. Gene number: GV: 562, MII: 174. **g,** Box plot of the ratio of read counts between dPAS and pPAS isoforms per gene on the log2 scale determined from PAIso-seq1 data of human oocytes at different stages. Gene number: GV: 2,051, MI: 441, MII: 169. **h,** Box plot of the poly(A) tail length per isoform for pPAS and dPAS isoforms determined from PAIso-seq1 data of human oocytes at different stages. Gene number: GV: 2,051, MI: 441, MII: 169. For all the box plots, the “×” indicates mean value, the black horizontal bars show the median value, and the top and bottom of the box represent the 25^th^ and 75^th^ percentiles, respectively. Genes with at least 10 poly(A) tail-containing reads (tail length ≥ 1) for both dPAS and pPAS isoforms are included in the analysis. *P* value is calculated by Student’s *t*-test.

**Fig. S3.**
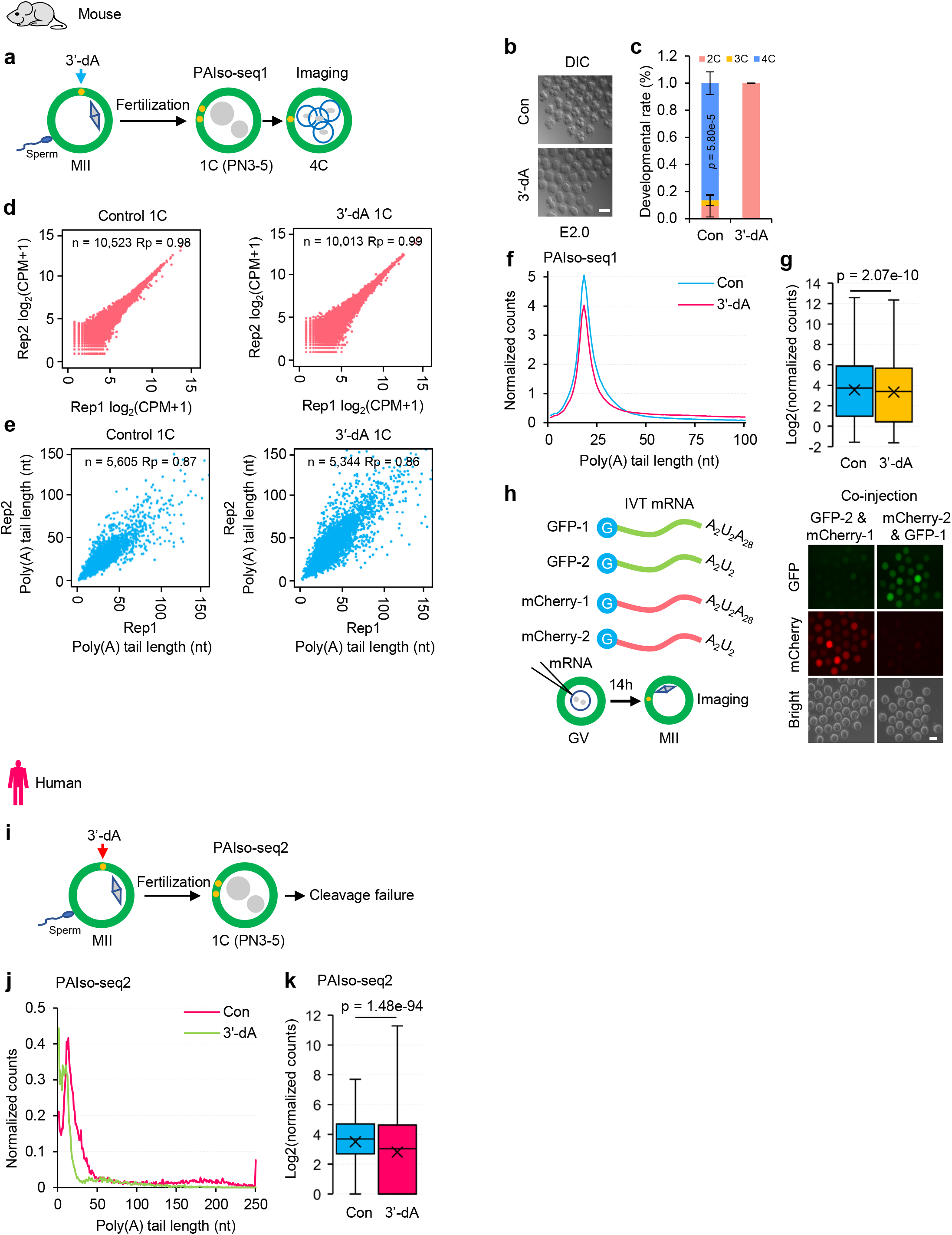
3’-dA inhibition of mRNA tail re-polyadenylation after fertilization in mice and humans. **a,** Illustration of 3’-dA treatment experiments in mouse oocytes. **b,** Morphology of control (Con) and 3’-dA-treated (3’-dA) mouse embryos at E2.0. Scale bar, 100 μm. **c,** Developmental rates of control and 3’-dA-treated mouse embryos examined at E2.0. Numbers of embryos analyzed are indicated (n = 85 for Con, n = 86 for 3’-dA). Error bars indicate the SEM from three replicates. **d,** Scatter plots showing the Pearson correlation of gene expression between two replicates for control and 3’-dA treated 1C embryos measured with PAIso-seq1. Each dot represents one gene. Pearson’s correlation coefficient (Rp) and number of genes included in the analysis are indicated at the top. **e,** Scatter plots showing the Pearson correlation of poly(A) tail length between two replicates for control and 3’-dA treated 1C embryos measured with PAIso-seq1. Each dot represents one gene. The poly(A) tail length for each gene is the geometric mean length of all transcripts with poly(A) tails at least 1 nt in length for the given gene. Genes with at least 20 reads in each sample were included in the analysis. Rp and number of genes included in the analysis are shown at the top. **f,** Histogram of poly(A) tail lengths of all transcripts in control or 3’-dA-treated 1C mouse embryos sequenced by PAIso-seq1. Transcripts with poly(A) tail lengths of 1–100 are shown in the histogram. **g,** Box plot of the normalized gene expression level of individual genes of control or 3’-dA-treated 1C mouse embryos sequenced by PAIso-seq1. **h,** Schematic of the translation reporter system based on *in vitro* transcription of transcripts for GFP or mCherry, which enables rapid assessment of the translation efficiency-altering effects of different types of poly(A) tails (Top left). Illustration of co-microinjection and *in vitro* culturing of mouse oocytes from the GV to the MII stage (Bottom left). Fluorescence microscopy image showing the translation of mRNA transcripts for GFP-Flag and mCherry-Flag reporter sequences bearing different types (i.e.: A2U2, A2U2A28) of poly(A) tails (right). Scale bar, 100 μm for all images. **i,** Illustration of 3’-dA treatment experiments in human oocytes. **j,** Histogram of poly(A) tail lengths of all transcripts in human control or 3’-dA-treated 1C embryos sequenced by PAIso-seq2. **k,** Box plot of the normalized gene expression levels of individual genes in human control or 3’-dA-treated 1C embryos sequenced by PAIso-seq2. The read counts are normalized by counts of reads mapped to protein-coding genes in the mitochondria genome if normalization is indicated. For all the box plots, the “×” indicates mean value, the black horizontal bars show the median value, and the top and bottom of the box represent the 25^th^ and 75^th^ percentiles, respectively. *P* value is calculated by Student’s *t*-test. Histograms (bin size = 1 nt) include transcripts with a poly(A) tail of at least 1 nt. For histograms with an X axis upper limit of 250, transcripts with a poly(A) tail length greater than 250 nt are included in the 250 nt bin.

## Notes

### Competing Interest Statement

The authors have declared no competing interest.

